# Estimating the Excitatory-Inhibitory Balance from Electrocorticography Data using Physics-Informed Neural Networks

**DOI:** 10.1101/2024.08.30.610583

**Authors:** Roberto C. Sotero, Jose M. Sanchez-Bornot

**Affiliations:** Department of Radiology, and Hotchkiss Brain Institute, University of Calgary, AB, Canada; Intelligent Systems Research Centre, Ulster University, Derry Londonderry, United Kingdom

**Keywords:** Electrocorticography, Physics-Informed Neural Network, Monkey, Anesthesia, neural mass model

## Abstract

Understanding the excitatory/inhibitory (E/I) balance in the brain is crucial for elucidating the neural mechanisms underlying various cognitive functions and states of consciousness. Mathematical models have provided significant insights into these mechanisms, but they often face challenges due to high dimensionality, noisy observation signals, and nonlinearities. In this paper, we introduce a novel methodology using Physics-Informed Neural Networks (PINNs) to estimate the E/I balance from electrocorticography (ECoG) data, effectively addressing these limitations. By integrating physical laws via a neural mass model with neural network training, our approach enhances parameter estimation accuracy and robustness. Our analysis reveals a significant reduction in long-range connections (LRCs) and excitatory short-range connections (SRCs) under anesthesia, alongside an increase in inhibitory SRCs, highlighting anesthesia’s role in modulating neural dynamics to induce unconsciousness. These findings not only corroborate existing theories on the neural mechanisms of anesthesia but also provide new insights into brain connectivity and its relationship with consciousness.

**CCS CONCEPTS:** Computing methodologies→Machine learning

**ACM Reference Format:** 

## 1 INTRODUCTION

The excitatory/inhibitory (E/I) balance refers to the dynamic regulation between excitatory neurotransmitters, such as glutamate, which promote neuronal firing, and inhibitory neurotransmitters, such as gamma-aminobutyric acid (GABA), which suppress neuronal activity [1]. E/I balance is fundamental to several brain functions, including sensory processing, learning, memory, and cognition. Precise regulation of E/I signals ensures that neurons communicate effectively without becoming overly excited or inhibited, which could lead to network instability or dysfunction. Disruptions in the E/I balance are implicated in various neurological and psychiatric disorders [2]. For instance, in epilepsy, an excess of excitatory activity or a deficit in inhibitory control can lead to uncontrolled neuronal firing, resulting in seizures [3]. In autism spectrum disorder (ASD), an altered E/I balance is thought to contribute to sensory processing abnormalities and cognitive deficits [2]. Schizophrenia is another condition where E/I imbalance is evident, with disrupted inhibitory signaling, particularly involving GABAergic interneurons, potentially underlying cognitive and perceptual disturbances [4]. The E/I balance is also profoundly affected during anesthesia. Anesthetic agents typically enhance inhibitory neurotransmission and/or diminish excitatory neurotransmission to induce a reversible loss of consciousness and sensation [5]. For example, ketamine, a commonly used anesthetic, acts as an NMDA receptor antagonist, leading to a decrease in excitatory signalling and modulation of cortical and subcortical neural activity [6]. Similarly, dexmedetomidine, another anesthetic, works as an alpha-2 adrenergic agonist, promoting inhibitory neurotransmission and further altering the E/I balance [7]. Understanding how different anesthetics manipulate the E/I balance is crucial for optimizing anesthetic protocols and improving patient safety.

Mathematical models have been pivotal in clarifying the complex interactions between excitatory and inhibitory neurons. These models provide frameworks to simulate and analyse the underlying mechanisms that maintain neural circuit stability and function. For instance, Vogels et al. [8] developed a model investigating how inhibitory plasticity contributes to maintaining the E/I balance. Their model incorporated inhibitory synaptic plasticity rules based on the timing of pre- and post-synaptic spikes. Their results demonstrated that inhibitory plasticity could dynamically adjust the strength of inhibitory synapses to stabilize network activity, preventing runaway excitation or excessive inhibition. Deneve and Machens [9] introduced a probabilistic framework to explore how the brain maintains the E/I balance to optimize sensory processing and information transmission. Their model suggested that the E/I balance is dynamically regulated to minimize prediction errors in sensory inputs, providing a theoretical basis for understanding how neural circuits maintain flexibility and robustness. Litwin-Kumar and Doiron [10] explored the role of structured connectivity in maintaining the E/I balance. Their model incorporated spatially organized connectivity patterns among excitatory and inhibitory neurons, reflecting cortical circuits’ anatomical and functional architecture. The results showed that structured connectivity could support a stable E/I balance by promoting localized synchrony and preventing widespread pathological activity.

While these theoretical works presented forward models that simulate the E/I balance in brain networks, further advancing our understanding requires solving the inverse problem. This involves estimating the parameters of these mathematical models using real EEG (electroencephalography) or ECoG (electrocorticography) data. By doing so, researchers can derive the specific excitatory and inhibitory dynamics that occur within actual brain networks, thereby providing a more precise and individualized understanding of neural function and dysfunction. Solving the inverse problem is crucial for translating theoretical models into practical applications, such as diagnosing and treating neurological disorders characterized by E/I imbalances. It allows for the refinement of models based on empirical data, leading to more accurate predictions and effective interventions [11]. However, these models often require extensive a priori knowledge and struggle with parameter estimation challenges arising from the noise and complexity inherent in EEG and ECoG data. When modelling extensive brain networks, the high dimensionality of such models [12] leads to considerable computational challenges in fitting them to empirical data.

The introduction of Physics-Informed Neural Networks (PINNs) [13] offers a promising solution to these issues. PINNs integrate physical laws described by partial differential equations (PDEs) with neural networks, enabling the incorporation of empirical data and theoretical models within a unified framework. This approach allows for more robust parameter estimation by leveraging the neural network’s ability to handle noisy and complex data [14].

In this paper, by embedding the governing equations of the E/I balance via a neural mass model (NMM) directly into the neural network training process, we develop a PINN that provides a powerful tool for accurately estimating model parameters from ECoG data, thus facilitating the translation of theoretical models into practical clinical applications.

## 2 MATERIALS AND METHODS

### 2.1 Macaque ECoG data

In this study we used ECoG data from a macaque monkey during rest and anesthesia (administered with a combination a combination of 5.00 mg of ketamine and 0.02 mg/kg of medetomidine) conditions [15]. The ECoG array consisted of 128 electrodes, with signals sampled at a rate of was 1000 Hz. For analysis, the time series for both rest and anesthesia conditions were divided into 300 segments, each lasting 1 second. These segments were filtered within the alpha band (5-12 Hz) and subsequently downsampled to a 100 Hz sampling rate to facilitate further analysis.

### 2.2 LSTM Networks

Long Short-Term Memory (LSTM) networks are a specialized type of Recurrent Neural Network (RNN) designed to effectively recognize patterns in sequences of data making them particularly suitable for tasks involving time series [16]. The fundamental concept behind an LSTM network is the cell state, which serves as a mechanism to preserve and convey relevant information throughout the sequence processing. LSTM units contain a set of gates that control the flow of information into and out of the cell state. The equations describing the LSTM unit at each time step *t* are:

- Forget Gate:

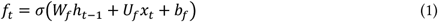
- Input Gate:

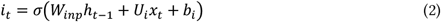

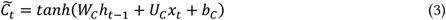
- Cell State Update

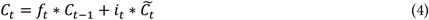
- Output Gate:

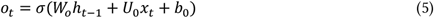

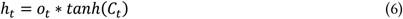

where *W* are the weights related to the input *x*_*t*_ for each of the gates and the cell state, *U* are the weights associated with the recurrent connections, *b* represents the bias term for each gate, *σ* is the sigmoid function and *tanh* is the hyperbolic tangent. *f*_*t*_, *i*_*t*_, *o*_*t*_, and 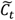 are the outputs of the forget gate, input gate, output gate, and the candidate cell state, respectively. *C*_*t*_ and *C*_*t*−1_ are the current and previous cell states, respectively. *h*_*t*_ and *h*_*t*−1_ are the current and previous hidden states, respectively. The asterisk (*) denotes element-wise multiplication.

### 2.3 Modelling the neural dynamics and the E/I balance

We describe the average postsynaptic potential (PSP), *y*_*i*_, generated on the pyramidal cells population and registered at electrode *i* by both local (short-range) and long-range activity, using the NMM framework [12]:

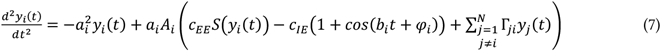

where *N* is the number of electrodes, and *a*_*i*_ and *A*_*i*_ represent the sum of the reciprocal of the time constants and the gain, respectively. The sigmoid function *S*(*x*) is given by the equation:

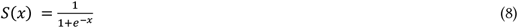

Short-range connections (SRCs) can be both excitatory (*c*_*EE*_) and inhibitory (*c*_*IE*_). The angular frequency *b*_*i*_ and phase *φ*_*i*_ are the parameters of the inhibitory PSP generated on the pyramidal cells population by local inhibitory interneurons. Long-range connections Γ_*ji*_ from other electrodes (*j*) are considered to be only excitatory[12].

We then model the E/I balance at electrode *i* as the ratio of excitatory connections to inhibitory connections:

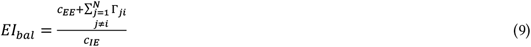

### 2.4 Parameter estimation within the PINN framework

In this paper, we individually fit the ECoG time series *y*_*i*_ for each electrode using the PINN framework, leveraging data from the other *N*-1 time series *y*_*j*_. Simultaneously, we estimate the biophysical parameters of the NMM, as detailed in Equation (7), including local parameters *a*_*i*_, *A*_*i*_, *c*_*EE*_, *c*_*IE*_, *b*_*i*_, *φ*_*i*_, and the *N*-1 LRCs Γ_*ji*_.

Figure 1 illustrates the PINN framework. We evaluate the PINN’s performance with a Mean Square Error (MSE) loss function that incorporates Data Loss and Physics Loss. Data Loss measures the differences between predicted and actual time series, while Physics Loss ensures compliance with the dynamics defined by the NMM’s differential equations. Our neural network, designed to estimate parameters that conform to observed data and physical laws, consists of two LSTM layers, with a dropout layer interposed to prevent overfitting, and a fully connected layer that predicts the PSP *y*_*i*_(*t*) at each timestep *t*. The number of LSTM hidden units is the same in the two layers and is denoted as *N*_*u*_; the dropout rate is *p*. The network inputs a matrix combining the time vector with the time series from all electrodes, excluding the target electrode *t*, and spans dimensions *N* × *T*, where *T* represents the ECoG series length.

**Figure 1:**
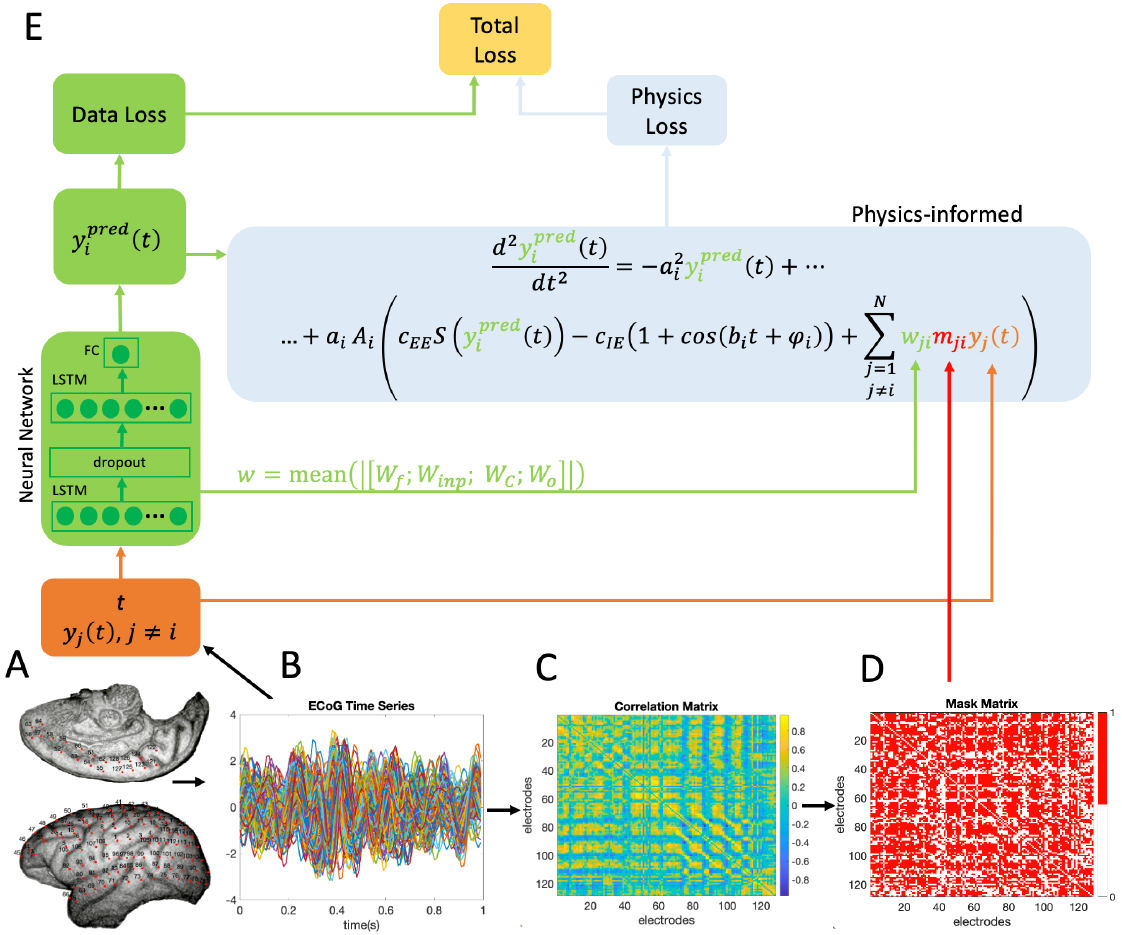
Framework of the Physics-Informed Neural Network (PINN) for estimating the neural mass model (NMM) parameters. A) Spatial distribution of the 128 ECoG electrodes. B) The 128 ECoG time series for a one-second interval. C) Correlation matrix constructed using pairwise Pearson correlation coefficients. D) Mask matrix with coefficients *m*_*ji*_ obtained by binarizing the correlation matrix. E) Model architecture: The neural network consists of two LSTM layers with a dropout layer between them. The inputs to the first LSTM layer are the time vector t and the time series from all electrodes (*y*_*j*_(*t*), *j* ≠ *i*). The network predicts *y*_*i*_(*t*) for each specific electrode. The Total Loss function is a composite measure consisting of Data Loss and Physics Loss. Data Loss evaluates the discrepancy between the predicted PSP and the observed ECoG data, while Physics Loss ensures compliance with the underlying NMM modeled by a second-order differential equation. This equation includes a term accounting for the influence from neuronal populations at other electrode locations, computed using the learned input weight coefficients *w*_*ji*_ from the first LSTM layer, multiplied by the binary mask matrix *m*_*ji*_.

The LSTM weights *W*_*f*_,*W*_*inp*_, *W*_*C*_, and *W*_*o*_ play a crucial role in processing the inputs. During training, we compute the vector *w* by averaging the absolute values of these input weight matrices across the LSTM units:

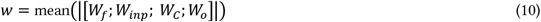

The first row, corresponding to the temporal feature, is discarded to preserve only the influences from the *N* − 1 electrode. The influence of each electrode *j* on electrode *i* is represented by *w*_*ji*_, obtained from *w*, and is used to calculate the LRCs Γ_*ji*_ as follows:

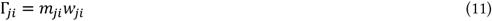

where *m*_*ji*_ is a mask matrix (Figure 1D) created by binarizing the correlation matrix (Figure 1C). This matrix is derived from the pairwise Pearson correlation coefficients between the ECoG time series (Fig 1B). The PINN was implemented using PyTorch [17] and underwent training for up to 1000 epochs. We employed the Adam optimizer [18] with the learning rate *α* treated as a hyperparameter. To optimize the model four hyperparameters— number of hidden units, dropout rate, learning rate, and L2 regularization coefficient—we utilized Bayesian optimization via the ‘hyperopt’ library [19].

## 3 RESULTS

For each condition (rest and anesthesia) we have 300 one-second ECoG segments. For each segment, we ran the PINN analysis for each electrode individually and fit the corresponding ECoG time series *y*_*i*_ using as input the time vector t and the *N*-1 time series from the other electrodes. Thus, for each electrode we estimated four hyperparameters (number of hidden units, dropout rate, learning rate, and L2 regularization coefficient), six local NMM parameters (*a*_*i*_, *A*_*i*_, *c*_*EE*_, *c*_*IE*_, *b*_*i*_, *φ*_*i*_), and the *N*-1 LRCs Γ_*ji*_. We then computed the E/I balance using Equation (9).

Figure 2 presents a detailed comparison of the estimated SRCs and LRCs and the computed E/I balance under resting and anesthesia conditions. Plot (A) shows that the LRCs (Γ_*ji*_) are significantly higher during rest compared to anesthesia, indicating a substantial reduction in long-range connectivity under anesthesia. This finding is consistent with previous studies that report a decrease in global brain connectivity during states of reduced consciousness such as anesthesia [20]. The significant drop in LRCs under anesthesia highlights the profound effect of anesthetic agents on the brain’s ability to maintain long-range neural communication. Plot (B) illustrates that the excitatory SRCs (*c*_*EE*_) are higher during rest, reflecting stronger local excitatory interactions in the resting state. In contrast, plot (C) reveals that inhibitory SRCs (*c*_*IE*_) are notably increased under anesthesia, indicating an enhancement of local inhibitory mechanisms. This shift towards inhibition is a well-documented phenomenon in the anesthesia literature, where increased inhibitory signaling is associated with the suppression of cortical activity and the induction of unconsciousness [21]. The differential modulation of excitatory and inhibitory SRCs underpins the overall changes in neural dynamics induced by anesthetic agents.

**Figure 2:**
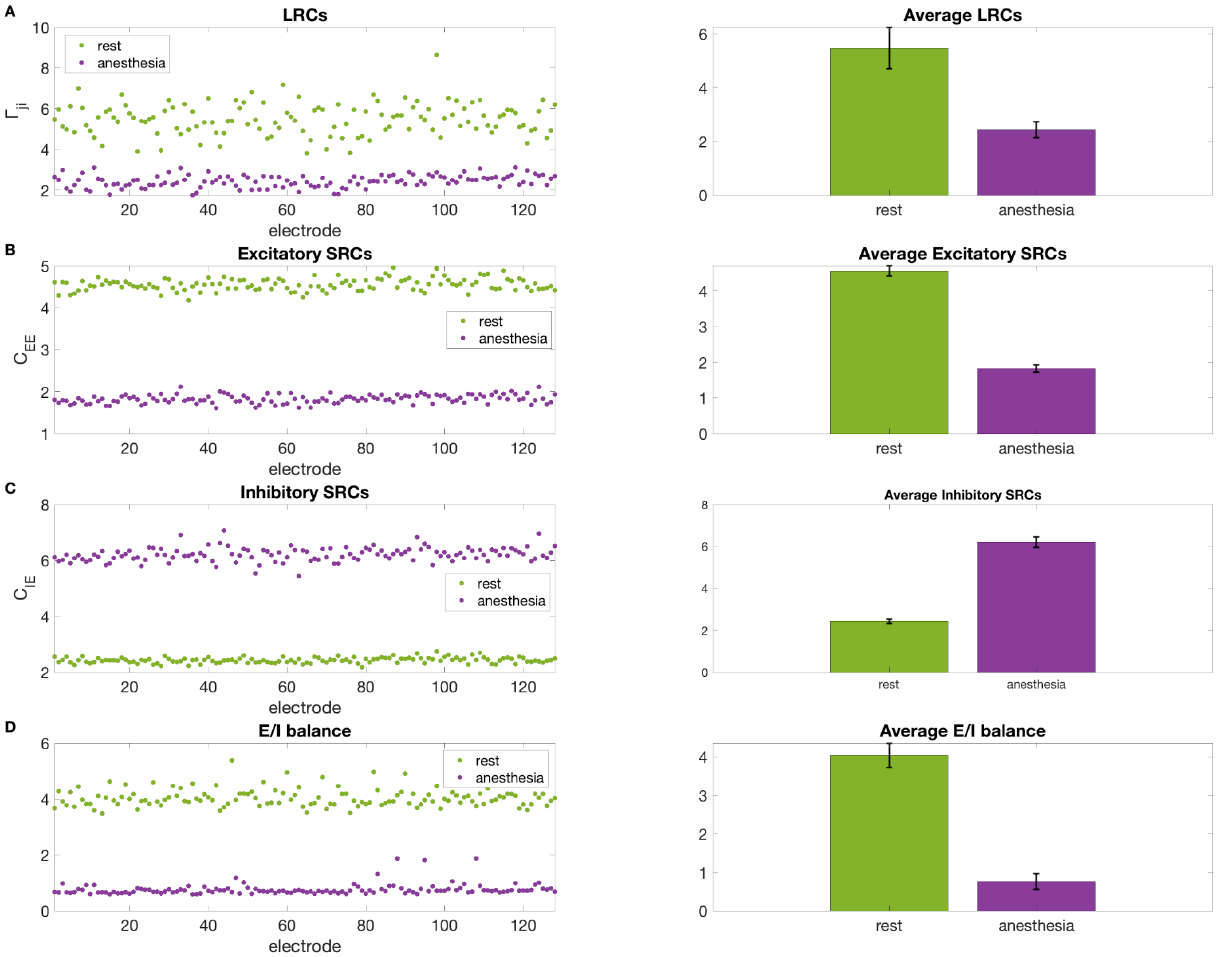
Analysis of the Excitatory/Inhibitory (E/I) balance under Rest and Anesthesia Conditions. A) LRCs (Γ_*ji*_) across 128 electrodes for rest and anesthesia conditions. The right panel shows the average LRCs with standard deviation. B) Excitatory SRCs (*c*_*EE*_) across 128 electrodes for rest and anesthesia conditions. The right panel shows the average excitatory SRCs with standard deviation. C) Inhibitory SRCs (*c*_*IE*_) across 128 electrodes for rest and anesthesia conditions. The right panel shows the average inhibitory SRCs with standard deviation. D) E/I balance across 128 electrodes for rest and anesthesia conditions. The right panel shows the average E/I balance with standard deviation. The results in plots A to D represent the average across 300 one-second intervals of the estimated parameters using the PINN framework detailed in Figure 1.

The focus of this paper is the E/I balance depicted in plot (D). During rest, the E/I balance is skewed towards excitation, with significantly higher values than those observed under anesthesia, which shows a dominance of inhibition. This observation is critical, as the E/I balance is a fundamental aspect of neural function, influencing the brain’s ability to process information and maintain cognitive functions. The shift towards inhibitory dominance under anesthesia aligns with the hypothesis that anesthesia induces unconsciousness by disrupting the normal E/I balance, thereby dampening neural excitability and reducing the overall level of cortical activity [5]. The bar plots on the right side of each panel summarize these findings, highlighting the substantial difference in average connectivity measures between the two conditions, with standard deviations indicating the variability across electrodes. The observed variability suggests that while the overall trend is towards reduced connectivity and increased inhibition under anesthesia, there are electrode-specific differences that may reflect the heterogeneous effects of anesthetic agents across different brain regions.

## 4 CONCLUSSIONS

We presented a novel approach utilizing PINNs for estimating the E/I balance within the brain from ECoG data. This approach addresses traditional neural mass modelling limitations such as nonlinearity, the large number of parameters to be estimated, noisy observation data, and high dimensionality. Our findings demonstrated a significant reduction in LRCs and excitatory SRCs, coupled with an increase in inhibitory SRCs. These changes underscore the role of anesthesia in modulating neural dynamics to induce unconsciousness.

Our results not only corroborate existing theories on the neural mechanisms of anesthesia but also offer valuable insights into the broader understanding of brain connectivity and its relationship with consciousness. The observed alterations in the E/I balance under anesthesia highlight the impact of anesthetic agents on neural connectivity, providing a clearer picture of how these agents influence brain function. Future studies should further explore the electrode-specific effects and investigate the underlying molecular and cellular mechanisms that contribute to these observed changes in connectivity and E/I balance under anesthesia. By doing so, we can enhance our understanding of the neural substrates of consciousness and the mechanisms through which anesthesia exerts its effects, ultimately improving clinical practices and patient outcomes.

## ACKNOWLEDGMENTS

This work was supported by grant RGPIN-2022-03042 from Natural Sciences and Engineering Council of Canada. The authors are grateful for access to the Tier 2 High-Performance Computing resources provided by the Northern Ireland High Performance Computing (NI-HPC) facility funded by the Engineering and Physical Sciences Research Council (EPSRC), Grant No. EP/T022175/1.

